# First evidence of Clavicipitaceous fungus attacking a vertebrate

**DOI:** 10.1101/2025.03.01.641002

**Authors:** Tsuyoshi Hosoya, Tomotaka Sato, Sho Kadekaru, Yumi Une

## Abstract

*Metapochonia bulbillosa* (*Clavicipitaceae*) was isolated as a causative agent of a new frog mycosis of *Pelophylax porosus brevipodus*, one of the endangered frogs in Japan. Analysis of spontaneous outbreaks and infection experiments confirmed that the fungus can lethally affect *Pelophylax porosus brevipodus*, as well a**s** *Dryophytes japonicus*. This is the first evidence that a species of *Clavicipitaceae*, originally known as a plant pathogen, also affects vertebrates.

## Introduction

Fungi interacts variously with other organisms-both plants, animals, and even fungi themselves through symbiosis to parasitism. Some parasitic interaction is highly destructive to their host, which even leads to the extinction of the hosts (Fisher et al. 2012). In clinical view, fungal infection for vertebrates are particularly important because of the risk for zoonoses that brings damages to both humans and animals (Bidaisee and Mcpherson 2014; Rahman et al. 2020; Rupasinghe et al. 2022). In most cases, the carrier animal are warm-blooded, while case reports of cold-blooded animals carrier are limited. However, in the recent years more diseases in cold-blooded animals are known such as snake fungal disease (SFD) and frog chytrids (Fisher et al. 2012).

Frogs are naturally distributed in all over the world, and one of the most popular cold-blooded animals as pets. Few reports on frogs are known. Since the discovery of chytridiomycosis caused by *Batrachochytrium dendrobatidis* Longcore, Pessier & D.K. Nichols (Longcore et al. 1999), fungal frog diseases are drawing public attention.

During May to July in 2023, abnormal ratio of serial death of *Pelophylax porosus brevipodus*, listed as Endangered IB in Japan (Ministry of the Environment, 2020), kept in an exhibition facility for ex situ conservation occurred in Japan. The samples were pathologically examined and diagnosed as mycosis. In this paper, we report the identification of the fungus and its infectivity and virulence to the frogs as demonstrated by experimental infection. This is the first evidence that Clavicipitaceous fungus, originally known as a plant pathogen, affects vertebrates.

## Materials and Methods

### Tissue samples

Thirty-three individuals excluding decomposed or mummified, were fixed in a 10% neutral buffered formalin solution for pathological examination, and fresh samples were taken for pathogen examination (results of epidemiological and pathological tests are being prepared for separate submission). All the liver tissues were frozen immediately after excised and kept at - 20 C until isolation. The liver tissues were thawed in room temperature, and each tissue particle was inoculated onto corn meal agar (CMA, Nissui, Tokyo) plate and incubated at room temperature (around 25 C) for 5 to 7 days.

### Isolation

Isolation of the fungi was carried out using a sterile needle by touching the spore drop formed at the tip of the mycelium growing out from the tissue. The isolates were grown on potato dextrose agar plate (PDA, Shimadzu, Tokyo). Isolation was attempted for multiple times for fungal colonies occurring from a single tissue. The plates were incubated at room temperature for about one week. After confirming the obtained colony appearances derived from a tissue particle were identical, one representative isolates were transferred to PDA slant. This procedure was repeated for 20 individuals. Representative isolates were deposited to NITE National Bioresource Center (NBRC). The same procedure was applied for livers excised from inoculated patient animals in inoculation experiment. For comparative purpose, *Metapochonia bulbillosa* JCM 18594 isolated from soil under *Acer mono* in Nagano Prefecture, Japan (Nonaka et al. 2013) was purchased from Japan Culture Collection (JCM).

### Morphological examination of the isolates

To observe morphology, PDA or CMA plate was inoculated with each obtained isolate and incubated at room temperature for about a week. Spore production structure was examined morphologically under light microscopy using Olympus BX-51.

### Molecular identification

For molecular identification, a 1 ml of 2% malt extract both, composed of 20 g of malt extract (Gibco, USA), 1 L of distilled water, was inoculated with the isolate, and incubated for about a week. The mycelium was transferred to a 2 ml round bottom Eppendorf tube, and freeze dried.

For DNA extraction from obtained isolates, polymerase chain reaction (PCR) amplification, and sequencing reaction with primer pair ITS1F/ITS4 (White *et al*., 1990) was used and the protocol in Itagaki, Nakamura & Hosoya (2019) was followed. Obtained sequences were assembled using the software ATGC version 7.0.3 (Genetyx, Tokyo, Japan) and BLAST searched in NCBI database (https://blast.ncbi.nlm.nih.gov/Blast.cgi).

### Infection experiment

To determine the infectivity and pathogenicity of frog-derived *Metapochonia bulbillosa* to frogs, infection experiments were carried out. Because *P. porosus brevipodus* is an endangered species, the experiment employed the surrogate of 12 wild-caught *Dryophytes japonicus* (Hylidae) using methods approved by the Certification number 24-A018 in Animal Experimental Committee of NAS Laboratory, Ltd., Japan. Since suspected fungal isolates were homogeneous, a representative isolate (FC-9885, deposited as NBRC 116752) and JCM 18594 were used for inoculation for comparative purpose. For control, uninoculated phosphate-buffered saline (PBS, pH 7.4) was used.

To prepare inocula, sterile water was poured to the slant and vigorously shaken by Vortex to release conidia. The suspension was collected and conidia density was checked with Thoma’s hemocytometer, and adjusted to 2 x 10^5^ conidia/ml. A 0.5 ml of the prepared suspension or control media was administrated by intra peritoneum (i. p.) or per os (p. o.)

After inoculation, frogs were kept in individual plastic cases and fed with bred crickets as diets. In the following 14 days of observation, any frogs that survived were sacrificed by hyperanaesthesia and fully autopsied. All organs, including dead frogs donated for pathological examination were fixed in 10% neutral buffered formalin solution. The other part of the liver tissues were provided for isolation of the fungus as previously described.

The recovered isolates were kept in PDA slants, and examined morphologically as previously described. For suspectable colonies, a part of mycelium was collected in the round bottom Eppendorf tube containing beads for molecular identification. Extraction of DNA, PCR, sequencing was carried out as previously described.

## Results

### Fungal isolates from frogs that died of natural causes

The materials were brought to the laboratory in two lots. From the liver samples obtained in the first lot, three out of four samples (F2-2, 4, 5 out of F2-2 to F2-5) showed some fungal elements. These fungal elements resulted in 5 isolates. All the second lot samples (W-1 to W-15) revealed fungal elements, and 13 isolates were successfully obtained from 13 samples. Aside from the possible contaminants, fungal isolates with the following characteristics recurred from the majority of the samples (Table 1).

**Table 1.**
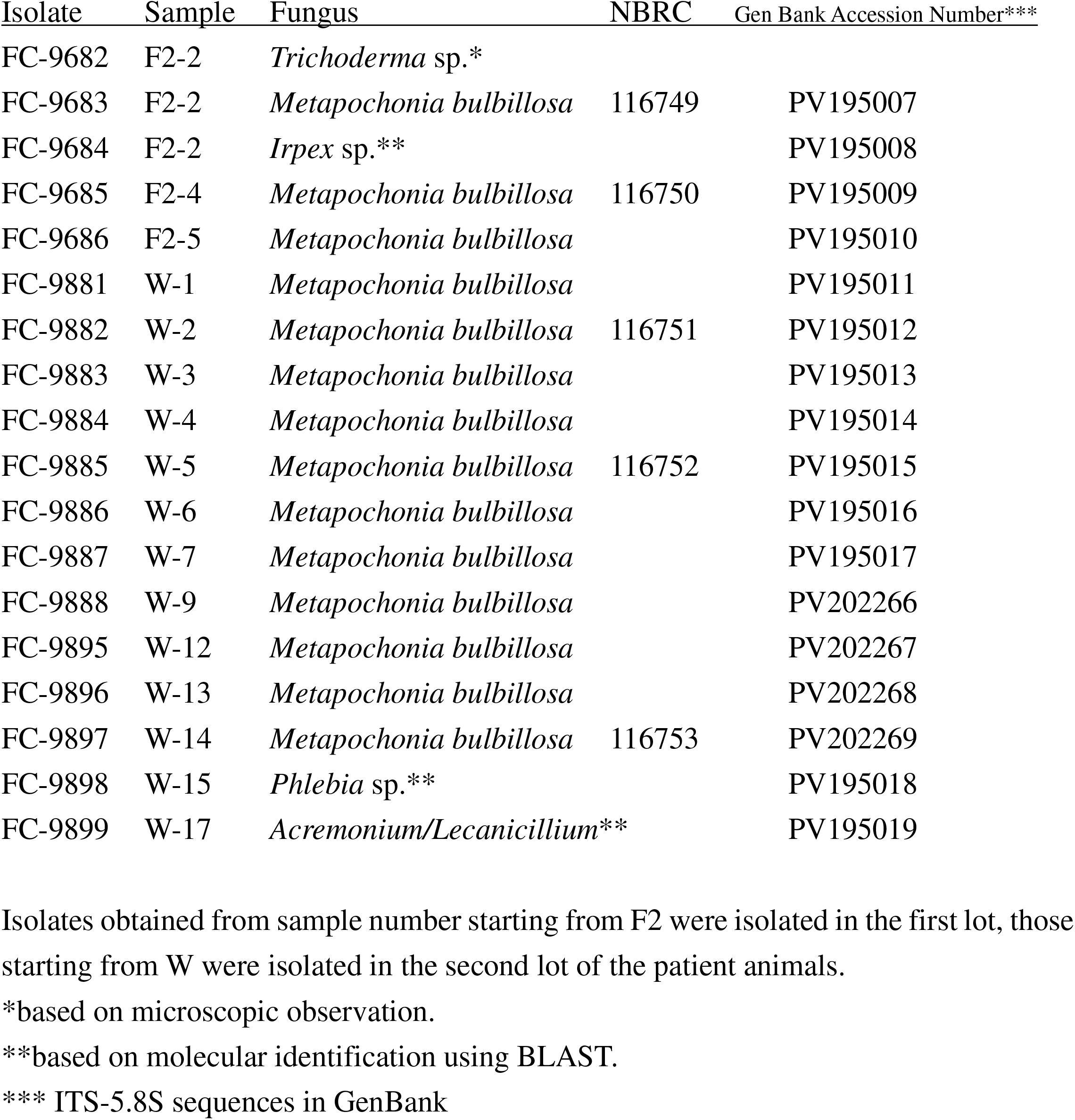
List of isolates obtained from liver of infected *Pelophylax porosus brevipodus*.

| Isolate | Sample | Fungus | NBRC | Gen Bank Accession Number*** |
| --- | --- | --- | --- | --- |
| FC-9682 | F2-2 | *Trichoderma* sp.* |  |  |
| FC-9683 | F2-2 | *Metapochonia bulbillosa* | 116749 | PV195007 |
| FC-9684 | F2-2 | *Irpex* sp.** |  | PV195008 |
| FC-9685 | F2-4 | *Metapochonia bulbillosa* | 116750 | PV195009 |
| FC-9686 | F2-5 | *Metapochonia bulbillosa* |  | PV195010 |
| FC-9881 | W-1 | *Metapochonia bulbillosa* |  | PV195011 |
| FC-9882 | W-2 | *Metapochonia bulbillosa* | 116751 | PV195012 |
| FC-9883 | W-3 | *Metapochonia bulbillosa* |  | PV195013 |
| FC-9884 | W-4 | *Metapochonia bulbillosa* |  | PV195014 |
| FC-9885 | W-5 | *Metapochonia bulbillosa* | 116752 | PV195015 |
| FC-9886 | W-6 | *Metapochonia bulbillosa* |  | PV195016 |
| FC-9887 | W-7 | *Metapochonia bulbillosa* |  | PV195017 |
| FC-9888 | W-9 | *Metapochonia bulbillosa* |  | PV202266 |
| FC-9895 | W-12 | *Metapochonia bulbillosa* |  | PV202267 |
| FC-9896 | W-13 | *Metapochonia bulbillosa* |  | PV202268 |
| FC-9897 | W-14 | *Metapochonia bulbillosa* | 116753 | PV202269 |
| FC-9898 | W-15 | *Phlebia* sp.** |  | PV195018 |
| FC-9899 | W-17 | *Acremonium/Lecanicillium* ** |  | PV195019 |

Isolates obtained from sample number starting from F2 were isolated in the first lot, those starting from W were isolated in the second lot of the patient animals.
\*based on microscopic observation.
\*\*based on molecular identification using BLAST.
\*\*\* ITS-5.8S sequences in GenBank

Colony on PDA attaining a diameter of 3.5–4.0 cm at 23C in 9 days, floccose, white from the surface, smooth to wrinkled, white to pale from the reverse. Phialides simple, branched, or verticillate, occasionally borne on short stalks, acicular, 20–50 mm long, 1.5–2 mm wide at the base. Conidia 2–7 x 1.5–2 mm, produced in drops at the head of phialides, almost spherical, ellipsoid to elongate, occasionally falcate (Figs. 1 and 2).

**Fig. 1.**
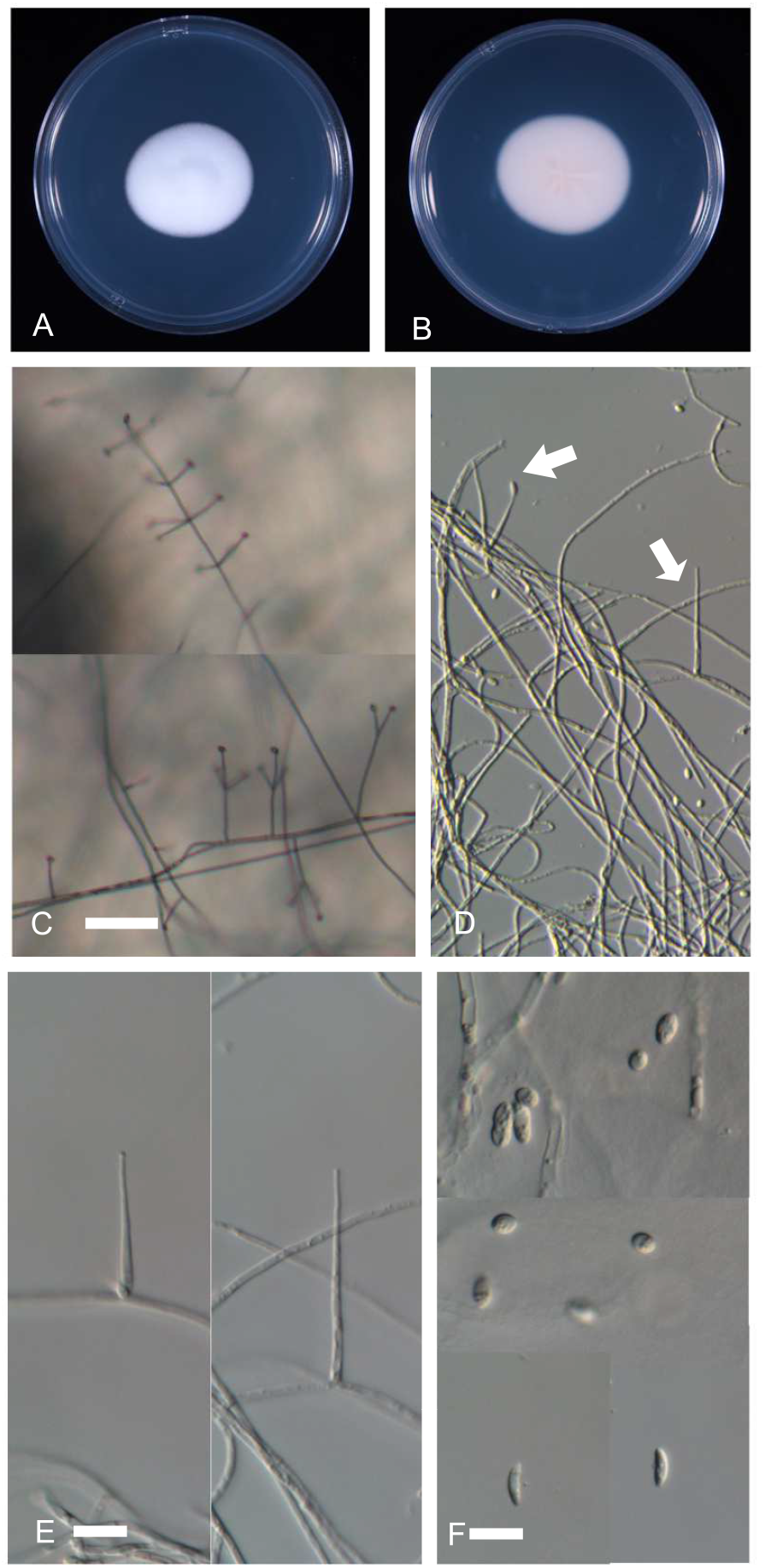
*Metapochonia bulbillosa* (FC-9885). (A) Colony on potato dextrose agar (23 C, 10 d), surface view. (B) Reverse view of the colony on PDA (C) Conidia producing structure on corn meal agar under low magnification. Verticillate phialides in upper photo, singly borne phialides and verticillate phialides on short stalks in lower photo. (D) Simple phialides. One at the left (arrow) have conidia borne at the tip. One at the right (arrow) is a acicular, simple phialide. (E) Simple phialides. (F) Conidia. Two at the bottom are falcate, characteristic to *M. bulbillosa*. Scale bars. C=50 μm, D=20 μm, E-F=10μm.

**Fig. 2.**
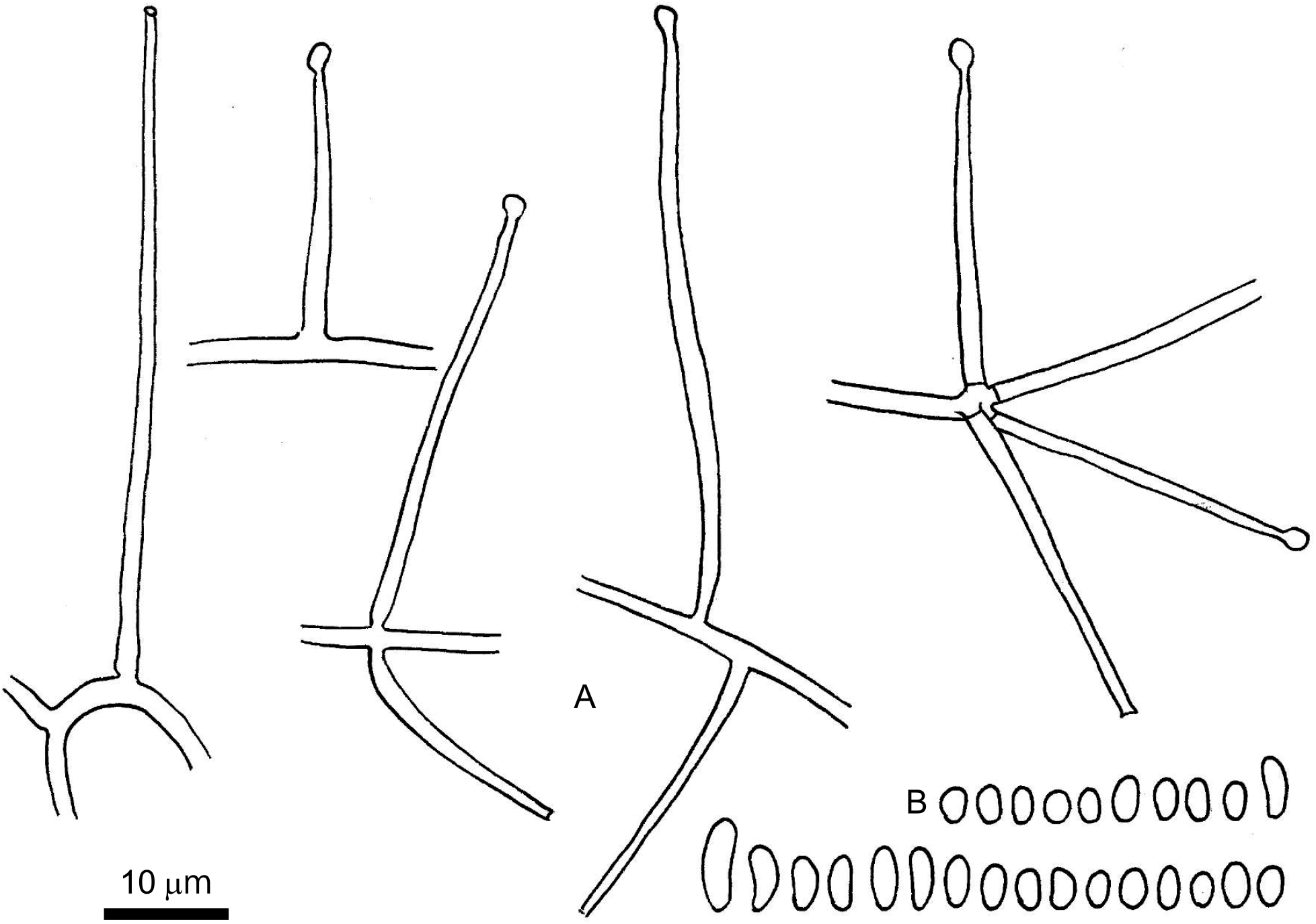
*Metapochonia bulbillosa* (FC-9885). Line drawings of the conidia producing structure produced on corn meal agar. (A) Various phialides born on hyphae. (B) Conidia. Some are falcate, characteristic to *M. bulbillosa*.

The morphology of these isolates, thought to be the putative pathogen, agreed with the description for *Metapochonia bulbillosa* (W. Gams & Malla) Kepler, S.A. Rehner & Humber (Gams 1971 ut *Verticillium bulbillosum* W. Gams, the basionym of *M. bulbillosa*), except for the lack of hyaline dictyochlamydospores. All the isolates obtained in the present study showed the asymmetrically curved conidia, the characteristics of the *M. bulbillosa* (Gams 1971).

The ITS-5.8S sequence obtained from both lots were identical, and suggested the serial death was caused by a single isolate or at least genetically close isolates. A representative sequence from FC-9885 was registered as (PV195015) in GenBank.

BLAST search based on ITS-5.8S also supported the identification. The sequence of FC- 9885 hit with *Pochonia bulbillosa* (a synonym of *Metapochonia bulbillosa*) with 100% percent identity and 100% query coverage.

### Pathological findings in dead frogs due to spontaneous infection with fungus

Subcutaneous edema, subcutaneous hemorrhage, skin ulceration, hemoperitoneum, and hemorrhage in the gastrointestinal tract were observed in the dead frogs. Histopathological examination revealed necrosis of hepatocytes and increased melanophages in the liver, septal thickening and necrosis of septal interstitial cells in the lungs, as well as an infiltration of macrophages containing pathogens coinciding with lesions in these organs (Fig. 3). The pathogen was spherical to oval in shape, approximately 1µm, with few hyphae observed. The pathogen stained red with PAS staining.

**Fig. 3.**
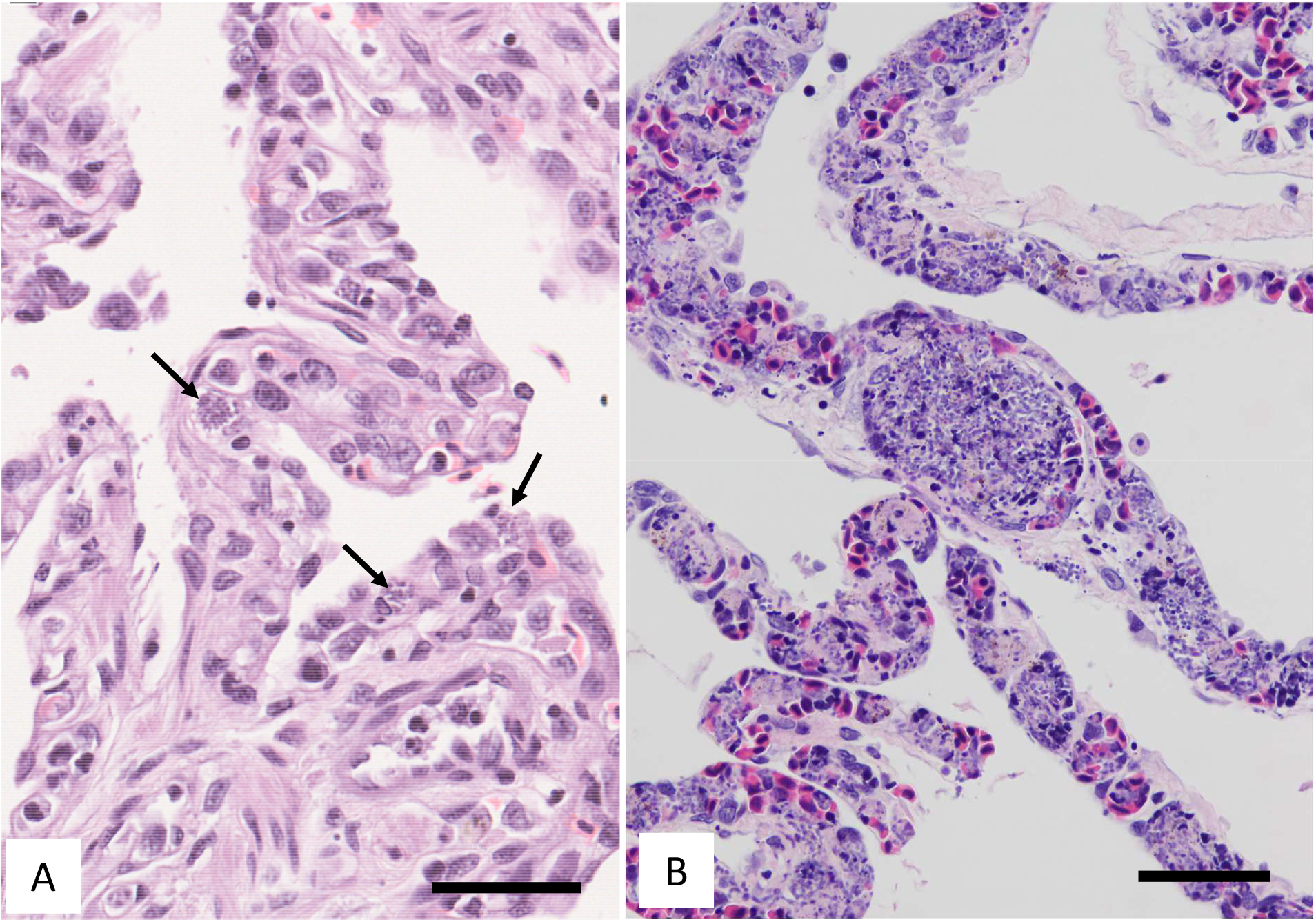
Pathological findings of lungs in frogs. (A)The fungus (arrows) is mainly present within the septa of lung of *Pelophylax porosus brevipodus*. (B) *Dryophytes japonicus* (DJ-1), inoculated with FC-9885. In the lung, a lot of fungi were observed within the septa. Scale bar =50 mm. Hematoxylin and eosin staining.

### Fungal isolates from frogs in inoculation experiments

Ten out of 12 liver samples revealed at least one fungus (Table 2). No fungus was detected in DJ_6 and 12 (control). While DJ_7 to 11 inoculated with JCM 18594 or PBS gave contaminants (*Arthrinium, Penicillium, Mucor*, and sterile hyphae), five (DJ_1 to 5) out of six samples (DJ_1 to 6) inoculated with FC-9885 gave white floccose hyphae as seen in *M. bulbillosa*. These were consisted of all three i. p. administration and two out of three p. o. administration.

**Table 2.** Results of infection experiment to *Dryophytes japonicus*.

| Sample | Inoculum | Administration* | Occurrence** | Initial<br>Weight (g) | Final<br>Weight (g) |
| --- | --- | --- | --- | --- | --- |
| DJ_1 | FC-9885 | i. p. | +(FC-9970, 9971) | 4.6 | 5.2 |
| DJ_2 | FC-9885 | i. p. | +(FC-9973) | 2.2 | 2.1 |
| DJ_3 | FC-9885 | i. p. | +(FC-9974) | 2.3 | 2.3 |
| DJ_4 | FC-9885 | p. o. | +(FC-9976) | 4.0 | 4.0 |
| DJ_5 | FC-9885 | p. o. | +(FC-9978) | 1.9 | 1.6 |
| DJ_6 | FC-9885 | p. o. | ND | 2.1 | 2.3 |
| DJ_7 | JCM 18594 | i. p. | - | 3.3 | 3.2 |
| DJ_8 | JCM 18594 | i. p. | - | 4.1 | 4.0 |
| DJ_9 | JCM 18594 | p. o. | - | 3.7 | 3.5 |
| DJ_10 | JCM 18594 | p. o. | - | 2.1 | 2.7 |
| DJ_11 | PBS (control) | p. o. | - | 3.5 | 3.7 |
| DJ_12 | PBS (control) | p. o. | ND | 2.7 | 2.8 |

\*i. p.: intra peritoneum; p. o.: per os; PBS. Phosphate-buffered saline (PBS, pH 7.4)
\*\*Occurrence shows if *Metapochonia bulbillosa* was recovered or not. +: present. -: absent. ND: no fungus detected. The isolate number indicated in the parenthesis represents those in the supplementary data 1.

The spore producing structure showed identical morphology with those of the original isolate. The obtained sequence from these isolates were identical with the sequence of the original isolate (Supplmentary data 1), and they were all identified as *M. bubillosa*. The inoculation experiment demonstrated no infection ability in JCM 18594, while FC-9885 demonstrated constant virulence in *D. japonica*.

### Pathological findings of frogs in inoculation experiments

During the observation period, all frogs showed no clinical signs of note. No remarkable weight change was observed during the inoculum experiment. On day 14 post-inoculation, one *D. japonicus* (DJ_1), which had been inoculated abdominally with FC-9885, died suddenly. Grossly, there were no significant lesions such as haemorrhage or ulceration of skin. Histopathologically, pathogens were only observed in two abdominally inoculated *D. japonicus* (DJ1 and DJ2). Pathogens showed identical morphology to that observed in naturally dead *Pelophylax porosus brevipodus*, with particularly large numbers of pathogens observed in the liver and lungs (Fig 3), and a high degree of necrosis of the liver cells.

## Discussion

### Inoculation experiments

From the above analyses, we concluded that *M. bubillosa* was the causal agent of this frog disease.

Whether the present symptom is a potential emerging disease or not is an intriguing question. The fact that the occurrence of this disease was highly contagious and rapidly expanded in the captured frogs suggests that *M. bulbillosa* potentially is a new agent of emerging disease. Although infection by *M. bubillosa* has not been confirmed in wild and captive frogs in Japan or overseas, this infection experiment suggests that it may at least be transmitted to and lethally affect wild native frogs in Japan. Since *M. bulbillosa* has already been isolated in Japan (Nonaka et al. 2013), it is not surprising that this is the discovery of overlooked pathogen in wild frog. However, we should also pay attention to the possibility that this is the emerging disease given that the disease was discovered in captured frogs.

### Mycological characteristics of *M. bulbillosa*

*Metapochonia bulbillosa* was originally described as a common soil fungus in the genus *Verticillium* [ut *Verticillium bulbillosum* (Kamyschko ex Barron & Onions) W. Gams], included in one of the largest section *Prostrata* (Gams 1971), characterized by both phialidic conidia with hyaline dictyochlamydospores, but the latter was not always accompanied (Zare et al. 2001). One of the remarkable characteristics of *M. bulbillosa* was the presence of falcate conidia at least partly asymmetrical along the longitudinal axis (Gams 1988). Dackman and Nordbring-Hertz (1985) reported that *V. bulbillosum* has rarely recorded as a parasite of cyst nematodes and ovicidal to eggs of certain nematode. Later, Zare et al. (2001) conducted a molecular study and adopted the genus *Pochonia* (asexul stage of *Cordyceps*) to accommodate this fungus. In the further treatment of *Codyceps*-related fungi, Kepler et al. (2014) disposed the species to a new genus *Metapochonia* based on the molecular phylogenetic analysis. Through these analyses, the taxaonomic position of this fungus became clearer, now placed in *Clavicipitaceae*, as a sister group of *Metarhizum* and *Pochonia*, both known to be predatory to animals. *Metarhizum* is insectivorous while *Pochonia* is known to be nematophagous. However, there has been no report of any parasitism to vertebrate. The infectivity and pathogenicity reported in the present study is the first example to show clear pathogenicity of member of *Clavicipitaceae* to vertebrates.

---

Supplementary data 1. Sequences obtained from recovered isolates from inoculated *Dryophytes japonicus*. The title of each sequence shows the isolate number (FC####) followed by GenBank accession number (PV######) followed by private sequence number (S#####). PCR carried out using ITS1F and ITS4.

